# The direct costs of overdiagnosed asthma in a longitudinal population-based study

**DOI:** 10.1101/409870

**Authors:** Bryan C. Ng, Mohsen Sadatsafavi, Abdollah Safari, J. Mark FitzGerald, Kate M. Johnson

**Affiliations:** Respiratory Evaluation Sciences Program, Collaboration for Outcomes Research and Evaluation, Faculty of Pharmaceutical Sciences, University of British Columbia, Vancouver, Canada; Institute for Heart and Lung Health, Department of Medicine, University of British Columbia, Vancouver, Canada

**Keywords:** Asthma, Costs, Misdiagnosis, Observational Studies, Overdiagnosis

## Abstract

**Objectives:** A current diagnosis of asthma cannot be objectively confirmed in many patients with physician-diagnosed asthma. Estimates of resource use in overdiagnosed cases of asthma are necessary to measure the burden of overdiagnosis and evaluate strategies to reduce this burden. We assessed the difference in asthma-related healthcare resource use between patients with a confirmed asthma diagnosis and those with asthma ruled out.

**Design:** Population-based prospective cohort study.

**Setting:** Participants were recruited through random-digit dialling of both landlines and mobile phones in BC, Canada.

**Participants:** We included 345 individuals ≥12 years of age with a self-reported physician diagnosis of asthma which was confirmed by a bronchodilator reversibility or methacholine challenge test at the end of the 12-month follow-up.

**Primary and secondary outcome measures:** Self-reported annual asthma-related direct healthcare costs (2017 Canadian dollars), outpatient physician visits, and medication use from the Canadian healthcare system perspective.

**Results:** Asthma was ruled out in 86 (24.9%) participants. Average annual asthma-related direct healthcare costs for participants with confirmed asthma were $497.9 (SD $677.9), and $307.7 (SD $424.1) for participants with asthma ruled out. In the adjusted analyses, a confirmed diagnosis was associated with higher direct healthcare costs (Relative Ratio [RR]=1.60, 95%CI 1.14-2.22), increased rate of specialist visits (RR=2.41, 95%CI 1.05-5.40) and reliever medication use (RR=1.62, 95%CI 1.09-2.35), but not primary care physician visits (p=0.10) or controller medication use (p=0.11).

**Conclusions:** A quarter of individuals with a physician diagnosis of asthma did not have asthma after objective re-evaluation. These participants still consumed a significant amount of asthma-related healthcare resources. The population-level economic burden of asthma overdiagnosis could be substantial.

**Strengths and limitations of this study:**

- Participants were recruited through random sampling of the general population in the province of British Columbia.
- Asthma diagnosis was confirmed or ruled out using sequential guideline-recommended objective airway tests.
- Healthcare resource use was self-reported, potential recall bias may have led to reduced accuracy.
- The study was unable to evaluate the indirect costs of overdiagnosis or the cost-savings from correcting the diagnosis.
- The generalizability of the results may be limited by regional differences in medical costs and practices.

## INTRODUCTION

Over 300 million people worldwide have been diagnosed with asthma.[1] Patients with asthma experience symptoms of wheezing, shortness of breath, chest tightness, and cough.[2] These symptoms, and periods of intensified disease activity referred to as exacerbations, impose a significant burden on healthcare resources and reduce patient quality of life.[3] A Canadian study estimated the excess direct medical costs of asthma at $1,058 (2013 Canadian dollars) per person-year.[4] The majority (74%) of asthma-attributed costs were due to medication use.[4]

Multiple evidence-based guidelines recommend confirming a diagnosis of asthma with objective testing for reversible airflow limitation or increased airway hyper responsiveness.[2,5] Despite these recommendations, previous studies suggest that in the community, asthma is diagnosed solely based on symptom history in over half of the cases.[6,7] The underuse of spirometry has been documented in Canada,[8] the United States,[9] and Europe.[10] A recent population-based study found that one in three patients with physician-diagnosed asthma did not meet the guideline-recommended spirometric criteria for asthma diagnosis and could have their medications safely stopped.[11] We refer to this group of patients as ‘overdiagnosed’.[12] These overdiagnosed patients are likely to be imposing costs on the healthcare system due to treatment for a condition that does not exist, and may be experiencing symptoms of an underlying illnesses that is not being treated.[11] By some estimates, there are over 785,000 overdiagnosed asthma patients in Canada alone.[13]

In response to these findings, some authors have called for population-based screening or case finding to re-evaluate previous diagnoses of asthma.[11,13] Assessing the value of these programs requires precise estimates of the burden of overdiagnosed asthma. To the best of our knowledge, estimates of the cost differences between overdiagnosed and confirmed cases of asthma currently do not exist. A previous study of the costs of overdiagnosed asthma was limited to assessing the potential asthma-related cost savings that a secondary screening program could provide.[13] Characterizing the patterns of healthcare resource use among patients in whom asthma can be ruled out can help identify opportunities for re-evaluation and inform initiatives to improve asthma diagnosis in the community.

We used a longitudinal population-based cohort of individuals with physician-diagnosed asthma to address this important evidence gap. Our primary objective was to compare total direct asthma-related healthcare costs in patients with a confirmed diagnosis of asthma versus patients in whom a diagnosis of asthma was ruled out using objective testing. Our secondary objectives were to characterize differences in healthcare resource use in terms of the 1) number of outpatient physician visits, and the 2) type and amount of asthma medication use.

## METHODS

### Study Design and Sample

We used longitudinal data from the Economic Burden of Asthma study (University of British Columbia Human Ethics #H10-01542), which has been previously described.[14,15] In summary, individuals with a self-reported physician diagnosis of asthma and at least one asthma-related healthcare encounter in the past 5 years were eligible. Participants were recruited through random-digit dialling of both landlines and mobile phones in the census subdivisions of Vancouver and Central Okanagan (populations of 603,502 and 179,839 in 2011, respectively) of BC, Canada.[16] Between 2010 and 2012, 618 participants were recruited, evaluated at baseline, and followed for 12 months with visits at 3-month intervals. We included 345 participants who were ≥12 years of age at baseline and successfully completed a bronchodilator reversibility or methacholine challenge test at the end of follow-up.

### Outcomes

Participants reported their asthma-related primary care and specialist physician visits, hospitalizations, emergency department visits, and current medication use at each study visit with a recall period of 3 months. The primary outcome was total asthma-related direct healthcare costs per patient over the one-year follow-up period. Total direct healthcare costs were comprised of all outpatient and inpatient encounters, and medication costs incurred by the patient. Cost categories were described by calculating the percentage of total average cost constituted by each category. The methods used for calculating costs are detailed elsewhere.[17,18] Briefly, costs for each patient were determined by multiplying self-reported resource use quantities by unit cost of each resource. We determined the average unit cost of an asthma-related physician visit (specialist vs. primary care) and hospitalization for the years 2008-2012 using International Classification of Diseases codes from provincial healthcare administrative data.[18] Medication unit costs were determined by linking Drug Identification Numbers of participant-reported asthma medications to the Provincial Drug Master Plan database.[19] Cost per dose was estimated using the lowest price equivalent of the medication. All costs were adjusted to 2017 Canadian Dollars[20] and the analysis was conducted from the perspective of the Canadian healthcare system.

Secondary outcomes were the number of asthma-related outpatient physician visits and use of asthma medications. We did not evaluate asthma-related emergency department visits or hospitalizations as a separate outcome due to the low frequency of these events (N=14). The number of outpatient physician visits over one-year of follow-up was assessed separately by physician type (primary care or specialist). Medication use was captured using the questionnaire shown in *Figure E1*, and medications were classified into controller or reliever using a reference list (*Table E1*). In general, controller medications are those with anti-inflammatory effects (namely inhaled corticosteroids and leukotriene receptor antagonists), while reliever medications are those that are used on as-needed basis for temporary symptom relief (namely short-acting beta agonists). Medication use was determined by calculating the Medication Possession Ratio (MPR) separately for controller and reliever medications. MPR represents the proportion of days in which medications were available to the participant over the follow-up period.[21]

### Exposure: Objective confirmation of asthma

Participants underwent an objective assessment of asthma at the final visit. The diagnostic algorithm for asthma is shown in *Figure 1* and consisted of both bronchodilator reversibility and methacholine challenge tests implemented in a stepwise fashion. Spirometry was performed by a trained technician using a regularly calibrated spirometer. Reversible airflow obstruction was defined as a ≥12% increase in Forced Expiratory Volume in 1 second (FEV_1_) 15 minutes after administration of 200mcg of salbutamol via pressurized metered dose inhaler and a spacer device.[22] Participants who did not meet the criteria for reversible airflow obstruction returned within one week to undergo a methacholine challenge test. Participants who did not meet the criteria for asthma diagnosis at the first methacholine challenge test had their controller medications tapered and discontinued by a respirologist before returning for a second methacholine challenge test. A diagnosis of asthma was ruled out if FEV_1_ decreased by <20% following the administration of 16mg/mL of methacholine[23] in both methacholine challenge tests. Participants had a ‘confirmed asthma diagnosis’ if they met the criteria for asthma at the bronchodilator reversibility test or either methacholine challenge test.

**Figure 1.**
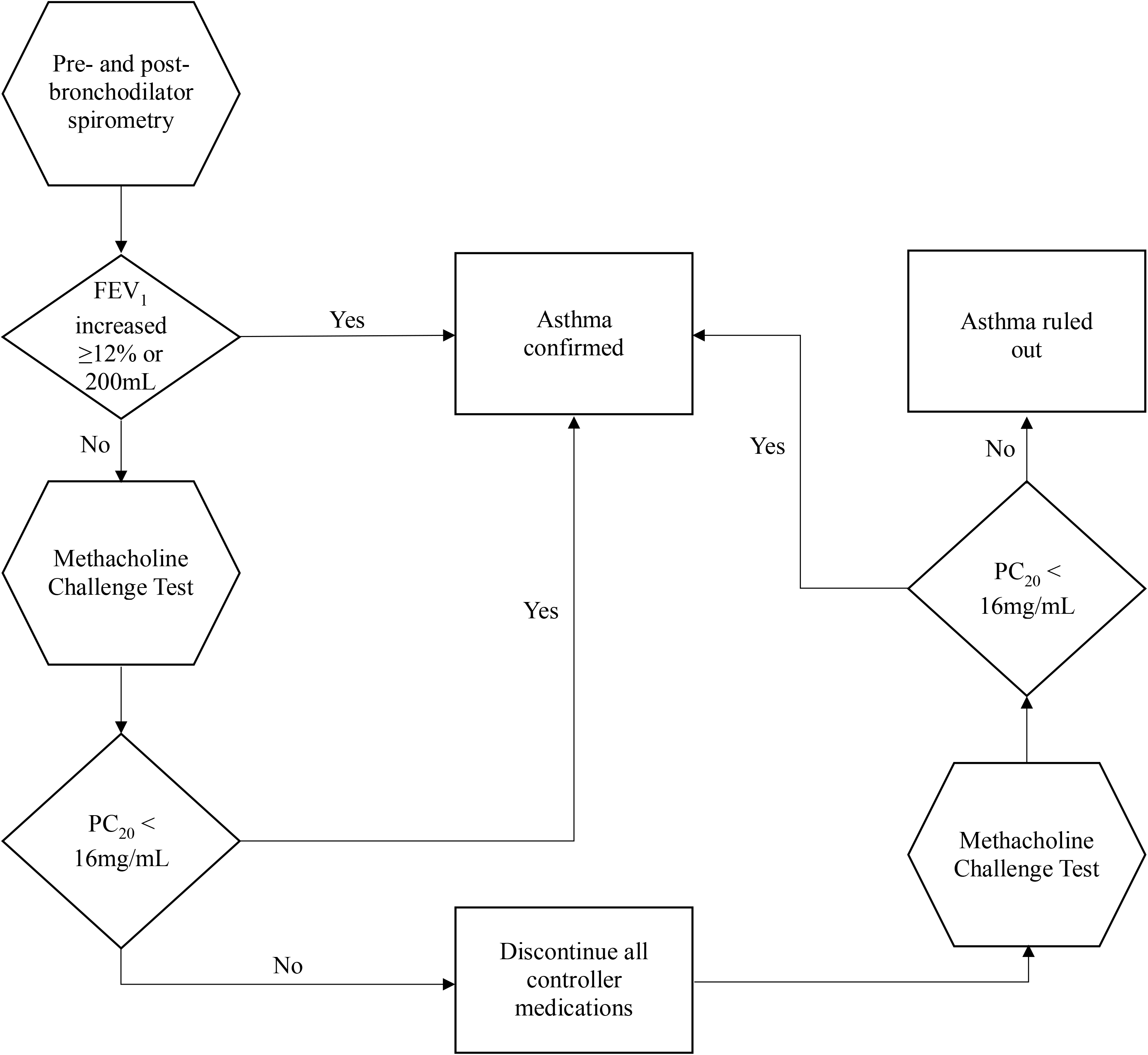
Algorithm for confirming or ruling out a diagnosis of asthma. FEV_1_; Forced Expiratory Volume in 1 second, PC_20_; Provocative concentration of methacholine needed to produce a 20% fall in FEV_1_ from baseline

### Statistical Analysis

All statistical analyses were performed in R version 3.5.0.[24] We considered a two-tailed p-value of <0.05 as statistically significant.

We constructed separate generalized linear regression models (negative binomial distribution, log link) for the primary and secondary outcomes. This resulted in five separate models for the following outcomes: annual asthma-related (1) direct healthcare costs, (2) number of primary care physician visits, (3) number of specialist physician visits, (4) controller MPR, and (5) reliever MPR. All models included the objective diagnosis of asthma (confirmed vs. ruled out) as the exposure. Costs and MPR values (as a percentage) were rounded to the nearest integer value. Models were adjusted for participant tobacco smoking history (ever smoked vs. never smoked), ethnicity (Caucasian vs. non-Caucasian), age, sex, education (postsecondary vs. no), income (annual household income ≥$70,000 vs. less), and third-party insurance coverage for medications (coverage vs. no), all self-reported at baseline with a 12-month recall period. The resulting regression coefficients were exponentiated to create effect estimates on the relative scale (rate ratio [RR]). Asthma severity was not adjusted for due to the high likelihood that there is a causal relationship between severe asthma and a confirmed diagnosis of asthma.[11]

## RESULTS

### Sample Characteristics

The cohort selection procedure is illustrated in *Figure 2*. From the Economic Burden of Asthma cohort (618 participants), we excluded 86 participants who were <12 years of age, 153 in whom a methacholine challenge was contraindicated (N=29), refused (N=112), or could not be completed for other reasons (N=14), and 34 participants who were lost to follow-up. The final study cohort included 345 participants who underwent an objective diagnostic test for asthma at the final visit (12 months after baseline). Their characteristics are shown in *Table 1*. 212 (61.4%) participants were female, and the mean age at baseline was 48.9 (standard deviation [SD] 17.8) years. A diagnosis of asthma was confirmed in 259 (75.1%) participants: 138 (53.3%) by bronchodilator reversibility test, 98 (37.8%) following the first methacholine challenge test, and 23 (8.9%) following the second methacholine challenge test. Asthma was ruled out in 86 (24.9%) participants following negative results on all bronchodilator reversibility and methacholine challenge tests.

**Figure 2.**
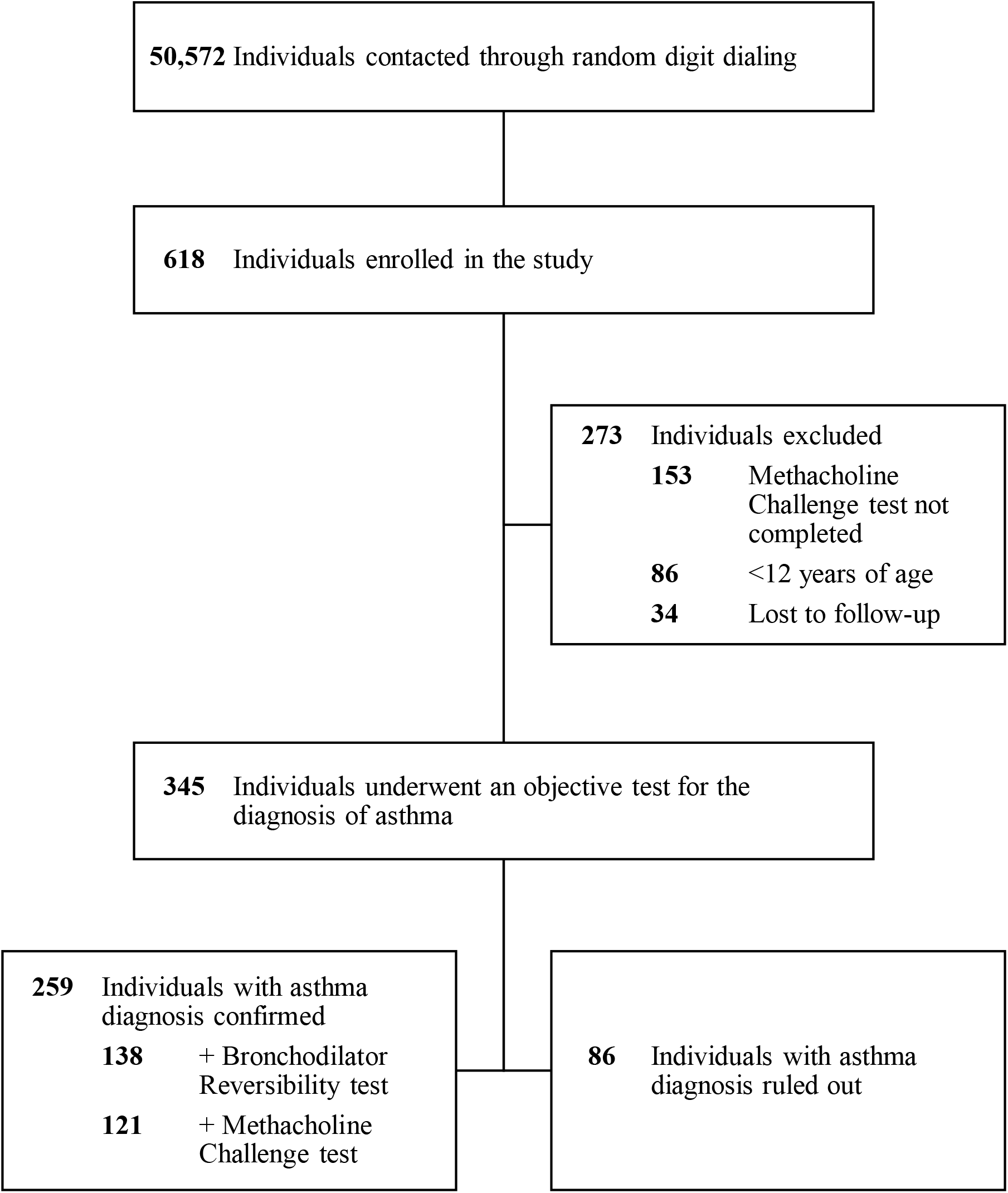
Sample selection procedure.

**Table 1.** Sample characteristics

| Characteristic | Total<br>(n = 345) | Asthma<br>confirmed<br>(n = 259) | Asthma<br>ruled out<br>(n = 86) | p-value§ |
| --- | --- | --- | --- | --- |
| *Variables evaluated at baseline* |  |  |  |  |
| Female,<br>n (%) | 212 (61.4) | 156 (60.2) | 56 (65.1) | 0.50 |
| Age,<br>mean (SD) | 48.9 (17.8) | 49.3 (18.1) | 47.7 (17.2) | 0.46 |
| Ever smoked (vs. never smoked),<br>n (%) | 93 (27.0) | 71 (27.4) | 22 (25.6) | 0.85 |
| Caucasian (vs. non-Caucasian),<br>n (%) | 267 (77.4) | 206 (79.5) | 61 (70.9) | 0.13 |
| High income (vs. low income) †,<br>n (%) | 183 (53.0) | 137 (52.9) | 46 (53.5) | >0.99 |
| Health insurance (full coverage<br>vs. not full coverage) ‡,<br>n (%) | 69 (20.0) | 52 (20.1) | 17 (19.8) | >0.99 |
| Higher education (postsecondary<br>vs. no postsecondary),<br>n (%) | 242 (70.1) | 178 (68.7) | 64 (74.4) | 0.39 |
| *Variables evaluated during follow-up* |  |  |  |  |
| Total direct healthcare costs,<br>mean (SD) | \$450.5<br>(\$629.2) | \$497.9<br>(\$677.9) | \$307.7<br>(\$424.1) | <0.01* |
| Primary care physician visits,<br>mean (SD) | 2.2 (4.1) | 2.1 (2.8) | 2.7 (6.6) | 0.85 |
| Specialist physician visits,<br>mean (SD) | 0.5 (1.9) | 0.6 (2.2) | 0.2 (0.8) | 0.02* |
| MPR of controller medications,<br>mean (SD) | 69.7% (74.4) | 74.9% (76.4) | 54.0% (65.8) | 0.02* |
| MPR of reliever medications,<br>mean (SD) | 28.4% (33.2) | 31.3% (34.3) | 19.5% (28.0) | <0.01* |

MPR: Medication Possession Ratio
\* Significant at 0.05 level
† High income defined as household income ≥ \$70,000
‡ Full coverage defined as all drug and physician services paid for through third-party coverage
§ p-values for the absolute difference between the 'asthma confirmed' and 'asthma ruled out'
groups were determined using a Student t-test for continuous variables and Chi-squared test for categorical variables

### Total Direct Healthcare Costs

Average annual direct healthcare costs over one year was $497.9 (SD $677.9) in participants with confirmed asthma, and $307.7 (SD $424.1) in those with asthma ruled out. There was a significant difference in average annual direct healthcare costs ($190.2) between exposure groups in the unadjusted analysis (p<0.01, *Table 1*). Average annual direct healthcare costs were 1.6 times (95%CI 1.14–2.22, p<0.01) higher in participants with confirmed asthma after adjustment for participant smoking history, ethnicity, age, sex, education, income, and insurance coverage (*Figure 3*). Medications comprised the greatest proportion of total costs in both correctly diagnosed and overdiagnosed individuals (56.4% and 47.6%, respectively, *Table 2*).

**Figure 3.**
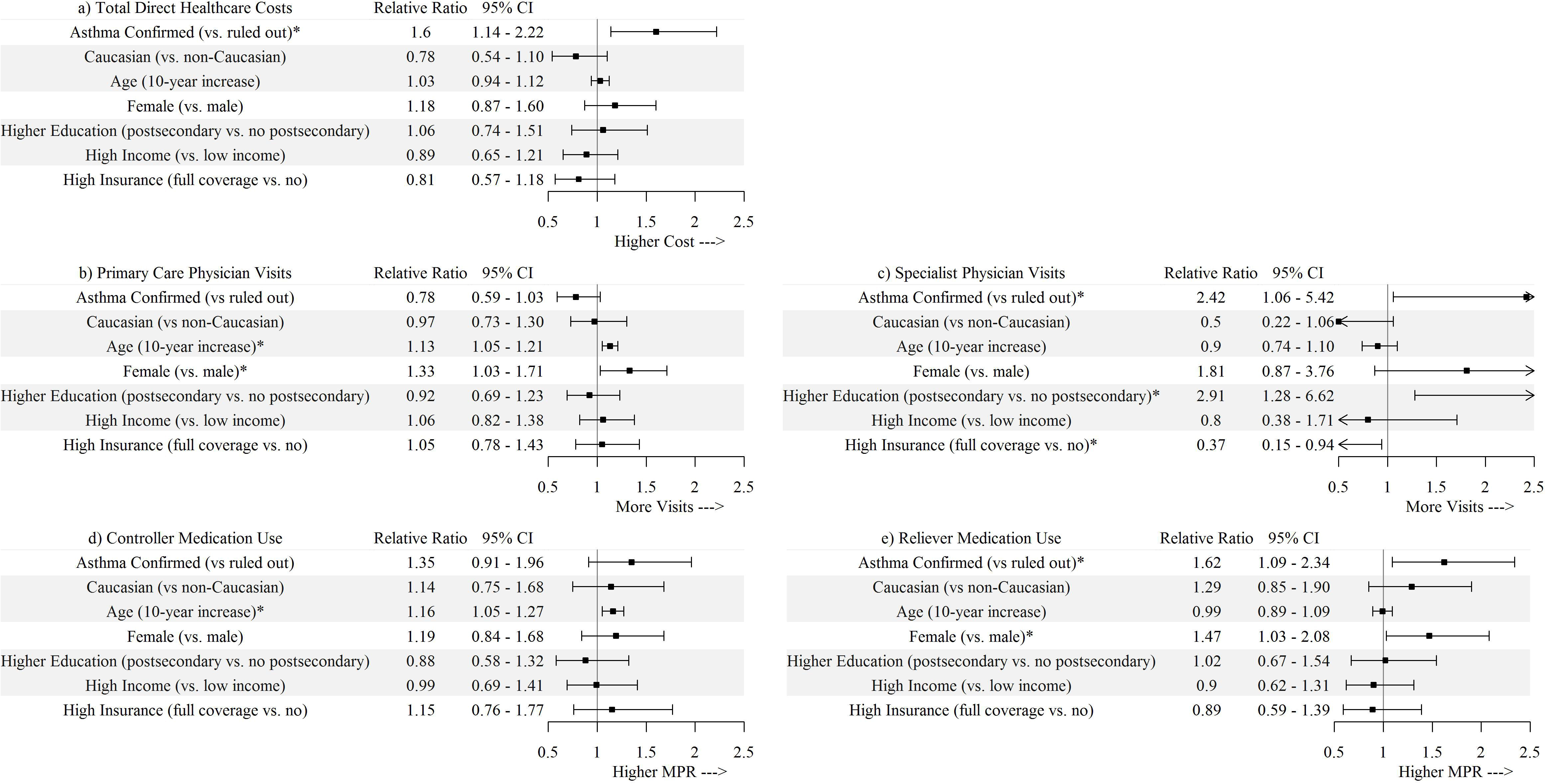
Relative Ratios for the adjusted differences in asthma-related a) total direct healthcare costs, b) primary care physician visits, c) specialist physician visits, d) controller medication use, and e) reliever medication use over one year of follow-up between patients with confirmed asthma and those with asthma ruled out. * Significant at 0.05 level

**Table 2.** Asthma-related healthcare cost categories over one year of follow-up.

| Cost Category | Cost per patient mean (SD) | Percentage of total cost per patient |
| --- | --- | --- |
| *Asthma ruled out* |  |  |
| Medications | \$146.39 (\$239.94) | 47.6% |
| Primary care physician visits | \$118.24 (\$292.64) | 38.4% |
| Specialist physician visits | \$27.87 (\$101.37) | 9.1% |
| Hospitalization | \$0 (\$0) | 0.0% |
| Emergency visits | \$15.23 (\$69.07) | 4.9% |
| TOTAL | \$307.70 (\$422.23) | 100.0% |
| *Asthma confirmed* |  |  |
| Medications | \$280.74 (\$404.17) | 56.4% |
| Primary care physician visits | \$90.98 (\$125.66) | 18.3% |
| Specialist physician visits | \$73.11 (\$257.61) | 14.7% |
| Hospitalization | \$39.24 (\$362.65) | 7.9% |
| Emergency visits | \$13.91 (\$72.07) | 2.8% |
| TOTAL | \$497.90 (\$676.89) | 100.0% |

### Outpatient Physician Visits

The total number of annual primary care physician visits was similar between participants with confirmed asthma and those with asthma ruled out (2.1 vs. 2.7 visits, p=0.85, *Table 1*). However, participants with confirmed asthma visited specialist physicians more often than those who had asthma ruled out (0.6 vs. 0.2 visits per year, p=0.02). The adjusted analysis also indicated no significant difference in the number of primary care physician visits between participants with confirmed asthma and those with asthma ruled out (p=0.10, *Figure 3*). Confirmed asthma was associated with 2.41 times (95%CI 1.05-5.40, p=0.03) more specialist visits than when an asthma diagnosis could be ruled out.

### Medication Usage

Participants with a confirmed diagnosis of asthma used controller medications for a greater proportion of follow-up time than those with asthma ruled out (MPR of 74.9% vs. 54.0%, p=0.02), and similarly for reliever medications (31.3% vs. 19.5%, p<0.01, *Table 1*). This difference persisted in the adjusted analysis for reliever medications but not for controller medications (p=0.11, *Figure 3*). The MPR for reliever medications was 1.62 times higher (95%CI 1.09-2.35, p=0.01) among participants with confirmed asthma than those with asthma ruled out.

## DISCUSSION

In this study, we used objective testing to confirm the diagnosis of asthma in a population-based sample of patients with a self-reported physician diagnosis of asthma. We found that asthma could be ruled out in 25% of cases after negative spirometry and two negative methacholine challenge tests. This proportion is in line with the 28-33% rate of overdiagnosis reported in previous Canadian studies.[11,13,25] We compared asthma-related direct healthcare costs between participants with confirmed asthma versus those with asthma ruled out. Although total direct costs were higher in participants with a confirmed diagnosis of asthma, the costs of overdiagnosed asthma remained substantial. The average direct asthma-related healthcare costs for a participant with overdiagnosed asthma was $308 over 12 months, which was $190 lower than for participants with a confirmed diagnosis. This difference in costs remained statistically significant after controlling for confounding variables that could affect both the exposure and outcomes but are likely not on the causal pathway between them.

Participants with overdiagnosed asthma visited specialists less frequently than those with confirmed asthma, but they visited primary care physicians as frequently. These participants may have scheduled a similar number of annual primary care visits for routine monitoring of ‘asthma’ activity or to fill new prescriptions. Conversely, participants with confirmed asthma may have had exacerbations of their asthma symptoms that required referral to a specialist physician to improve asthma control.[26] Across both groups, participants visited primary care physicians more frequently than specialist physicians; 79% of our sample visited a primary care physician at least once during follow-up. The high frequency of these visits, and their low cost compared to specialist consultations,[13] suggests that primary care visits may provide effective opportunities for re-evaluating previous asthma diagnoses.

On average, overdiagnosed participants possessed controller medications for over half of follow-up time, and reliever medications for 20 percent of follow-up time. Reliever medications are typically used as needed while controller medications are used daily.[2,5] In comparison to participants with confirmed asthma, and after adjusting for potential confounders, participants with asthma ruled out tended to use similar levels of controller medications and less reliever medications. This difference may be due to a lower symptom burden in overdiagnosed participants, which led to self-adjustment of reliever medication use.[27] Conversely, overdiagnosed individuals may have used controller medications as prescribed.

Although inhaled medications for asthma are generally safe, the use of asthma medications among overdiagnosed patients puts them at risk of net harm due to medication side effects.[28] These patients are also incurring additional healthcare expenditure without therapeutic benefit. Further, asthma medications may have masked symptoms of a serious underlying illness and resulted in a delay in the diagnosis and treatment of the correct disease. We did not evaluate the true underlying condition in this sample, but a similar study by Aaron et al.[11] found that two out of 213 patients had subglottic stenosis, which was treated as asthma for a number of years before the correct diagnosis was identified during the study. If other health conditions were responsible for the overdiagnosis of asthma, their costs have implications for the burden of overdiagnosed asthma. For example, if ruling out an asthma diagnosis results in the correct alternative diagnosis, the proper management of the underlying condition could confer further cost benefits.

To our knowledge, this is the first study to examine the costs associated with overdiagnosed asthma. Our findings have important population-level implications. Given an estimate of approximately 785,000 individuals with overdiagnosed asthma in Canada,[13] and an annual direct healthcare cost of $308 per patient, the estimated cost of overdiagnosed asthma in Canada is $242 million per year (2017 Canadian dollars). A previous study reported an average cost of $263 per patient (2009 Canadian dollars) for additional physician visits to reassess an asthma diagnosis. This resulted in an average lifetime cost savings of $351 per patient screened,[13] primarily due to the avoided costs of asthma medications. Our results suggest that additional screening to correct an overdiagnosis of asthma would save asthma-related costs in the first year, with savings compounding in subsequent years. Savings could be reallocated to the management of individuals with a true diagnosis of asthma, especially those with severe asthma who could pursue novel but expensive treatments, or to identify and treat the underlying diseases of overdiagnosed patients.[29] Given the prevalence and costs of overdiagnosis, our findings highlight the need for routine objective testing to confirm all new and existing diagnoses of asthma.[2,5]

There are several strengths to this study. We used a population-based random sample of patients with a physician diagnosis of asthma, meaning our results are likely to be representative of routine healthcare use in the general asthma population. We used longitudinal healthcare utilization data, which compared to cross-sectional data, likely provides a less-biased estimate of the average costs accrued by an individual. Finally, we evaluated airway reversibility using objective testing following the recommendations of international guidelines.[2,5] This allowed us to evaluate the costs of evidence-based best practices.

Our study has several limitations. We did not include indirect costs in our analysis, and we were therefore unable to consider the societal costs of overdiagnosis. Previous studies suggest that the cost of productivity loss in this sample is substantial.[15] Physician visits and medication use were self-reported with a recall period of 3 months. It is possible that recall bias reduced the accuracy of our measurements. Our algorithm for ruling out asthma involved one less methacholine challenge test than in some previous studies,[11,13,30] which makes it slightly less rigorous. It is possible that this led to a higher rate of overdiagnosis in our study, however, over 90% of diagnoses were confirmed by the second visit in previous studies.[13,25] We excluded participants in whom an objective diagnostic test was contraindicated due to asthma-related reasons, which may have resulted in lower representation of participants with more severe disease and therefore higher resource utilization. Finally, we only assessed asthma-related costs; we were unable to determine the costs of the true underlying condition or the benefit of correcting the diagnosis.

## CONCLUSIONS

In this population-based sample, one in four participants with a physician diagnosis of asthma had their diagnosis ruled out upon objective airflow reversibility testing. Patients with overdiagnosed asthma consumed a substantial amount of asthma-related healthcare resources, although less than those with confirmed asthma. The extent of overdiagnosed asthma in Canada and other countries, and its associated healthcare resource costs, suggests that the population-level burden of overdiagnosed asthma is high. Future studies should evaluate the cost-effectiveness of systematic screening or case detection initiatives for re-evaluating previous diagnoses of asthma.

## Acknowledgements

We would like to thank all members of the Economic Burden of Asthma study team. JMF and MS conceived, designed, and conducted the Economic Burden of Asthma study. MS formulated the current study idea and MS, KMJ and BCN designed the study. BCN performed all data analyses and wrote the first draft of the manuscript. AS and KMJ provided guidance on the statistical analysis. All authors critically commented on the manuscript and approved the final version. KMJ is the guarantor of the manuscript.

## ABBREVIATIONS

BC: British Columbia
CI: Confidence Interval
FEV_1_: Forced Expiratory Volume in 1 second
MPR: Medication Possession Ratio
SD: Standard Deviation

