## Supplemental Figure 1 for "The direct costs of overdiagnosed asthma in a longitudinal population-based study"

### CONFIDENTIAL

#### SECTION 3: Asthma Medications: This section will be completed by the interviewer.

1.1. During the past 3 months, have you taken any of the medications listed at your last visit for your breathing (also include medications for nasal congestion, reflux, allergies , and asthma exacerbation (s) such as prednisone and antibiotics)? ☐1.Yes ☐2.No ; If yes complete table below. If yes or no ask Q2.

1.2. During the past 3 months, have you taken any **new medications** listed for your breathing (including medications for nasal congestion reflux, allergies , and asthma exacerbation such as prednisone and antibiotics)? ☐1.Yes ☐2.No ; If yes complete table below.

If participant has not taken any medications to help their breathing, skip to Question 3.2.

| | | | | | | | |
|---|---|---|---|---|---|---|---|
| **Medication**<br>**Name** (not entered) | | | | | | | |
| *Medication Code (consult the medication name and code list for number)* | ___ ___ ___ | ___ ___ ___ | ___ ___ ___ | ___ ___ ___ | ___ ___ ___ | ___ ___ ___ | ___ ___ ___ |
| *Physician prescription Dose per use & frequency per day (e.g. 100mcg TID)* | | | | | | | |
| *Formulation (check one. If more than two formulations for one drug enter them as separate columns)* | Pills ☐ 1<br>Diskus ☐ 2<br>Inhaler ☐ 3<br>Nebulizer ☐ 4<br>Liquid ☐ 5<br>Nasal Spray ☐ 6<br>Injection ☐ 7<br>Other ☐ 8 | Pills ☐ 1<br>Diskus ☐ 2<br>Inhaler ☐ 3<br>Nebulizer ☐ 4<br>Liquid ☐ 5<br>Nasal Spray ☐ 6<br>Injection ☐ 7<br>Other ☐ 8 | Pills ☐ 1<br>Diskus ☐ 2<br>Inhaler ☐ 3<br>Nebulizer ☐ 4<br>Liquid ☐ 5<br>Nasal Spray ☐ 6<br>Injection ☐ 7<br>Other ☐ 8 | Pills ☐ 1<br>Diskus ☐ 2<br>Inhaler ☐ 3<br>Nebulizer ☐ 4<br>Liquid ☐ 5<br>Nasal Spray ☐ 6<br>Injection ☐ 7<br>Other ☐ 8 | Pills ☐ 1<br>Diskus ☐ 2<br>Inhaler ☐ 3<br>Nebulizer ☐ 4<br>Liquid ☐ 5<br>Nasal Spray ☐ 6<br>Injection ☐ 7<br>Other ☐ 8 | Pills ☐ 1<br>Diskus ☐ 2<br>Inhaler ☐ 3<br>Nebulizer ☐ 4<br>Liquid ☐ 5<br>Nasal Spray ☐ 6<br>Injection ☐ 7<br>Other ☐ 8 | Pills ☐ 1<br>Diskus ☐ 2<br>Inhaler ☐ 3<br>Nebulizer ☐ 4<br>Liquid ☐ 5<br>Nasal Spray ☐ 6<br>Injection ☐ 7<br>Other ☐ 8 |
| *When you are taking the medication, how many months since your last visit have you taken it?* | 0 ☐ 1<br>1 mth ☐ 2<br>2-3 mth ☐ 3<br>3-6 mth ☐ 4 | 0 ☐ 1<br>1 mth ☐ 2<br>2-3 mth ☐ 3<br>3-6 mth ☐ 4 | 0 ☐ 1<br>1 mth ☐ 2<br>2-3 mth ☐ 3<br>3-6 mth ☐ 4 | 0 ☐ 1<br>1 mth ☐ 2<br>2-3 mth ☐ 3<br>3-6 mth ☐ 4 | 0 ☐ 1<br>1 mth ☐ 2<br>2-3 mth ☐ 3<br>3-6 mth ☐ 4 | 0 ☐ 1<br>1 mth ☐ 2<br>2-3 mth ☐ 3<br>3-6 mth ☐ 4 | 0 ☐ 1<br>1 mth ☐ 2<br>2-3 mth ☐ 3<br>3-6 mth ☐ 4 |
| *When you are taking the medication, how many days a week do you take it?* | ☐☐ Days | ☐☐ days | ☐☐ days | ☐☐ days | ☐☐ days | ☐☐ days | ☐☐ Days |
| **In the past 3 months, is the medicine taken on most days, or just when yo have symptoms, or both?** | Most Days ☐ 1<br>Symptoms ☐ 2<br>Both ☐ 3 | Most Days ☐ 1<br>Symptoms ☐ 2<br>Both ☐ 3 | Most Days ☐ 1<br>Symptoms ☐ 2<br>Both ☐ 3 | Most Days ☐ 1<br>Symptoms ☐ 2<br>Both ☐ 3 | Most Days ☐ 1<br>Symptoms ☐ 2<br>Both ☐ 3 | Most Days ☐ 1<br>Symptoms ☐ 2<br>Both ☐ 3 | Most Days ☐ 1<br>Symptoms ☐ 2<br>Both ☐ 3 |
