## Supplemental Table 1 for "The direct costs of overdiagnosed asthma in a longitudinal population-based study"

**e-Table 1:** Asthma medication reference list.

| Medication categories | Active ingredient(s) | DIN |
| --- | --- | --- |
| Controller Medications | | |
| Inhaled corticosteroid (ICS) | Beclomethasone | 2242030, 2242029, 374407, 828521, 828548, 872334, 893633, 897353, 1949993, 1950002, 2079976, 2213710, 2213729, 2215039, 2215047, 2215055, 2216531 |
|  | Budesonide |  |
|  | Fluticasone |  |
| Long-acting beta- agonists (LABA) | Salmeterol | 2211742, 2214261, 2231129, 2136139, 2136147 |
|  | Formoterol | 2230898, 2237224, 2237225 |
| ICS and LABA in combination (ICS+LABA) | Budesonide, formoterol | 2245385, 2245386 |
|  | Fluticasone, salmeterol | 2240835, 2245126, 2245127, 2240836, 2240837 |
| Leukotriene receptor antagonists (LTRA) | Montelukast | 2247997, 2238217, 2243602, 2238216 |
|  | Zafirlukast | 2236606 |
| Anti-immunoglobulin E monoclonal antibody | Omalizumab | 2260565 |
| Inhaled mast cell stabilizers | Cromoglicic acid (cromolyn) | 2231431, 2231671, 2046113, 534609, 555649, 261238, 638641, 2049082, 2219468 |
| Theophylline | Choline theophyllinate | 346071, 405310, 441724, 441732, 451282, 458708, 458716, 476366, 476390, 476412, 503436, 511692, 536709, 565377, 589942, 589950, 792934 |
|  | Theophylline | 156701, 261203, 460982, 460990, 461008, 466409, 488070, 532223, 556742, 575151, 599905, 627410, 631698, 631701, 692689, 692697, 692700, 722065, 1926586, 1926594, 1926608, 1926616, 1926640, 1966219, 1966227, 1966235, 1966243, 1966251, 1966278, 1966286, 2014165, 2014181, 2230085, 2230086, 2230087 |
|  | Aminophylline | 14923, 178497, 497193, 497193, 497207, 582654, 582662, 868450, 2014270, 2014289 |
| Other corticosteroids | Cortisone | 280437, 16241, 16446, 16438 |
|  | Triamcinolone | 2194090, 15016, 15024, 2194082 |
|  | Prednisone | 610623, 598194, 550957, 312770, 252417, 210188, 868426, 868434, 868442, 21695, 232378, 607517, 508586, 156876, 271373, 271381 |
|  | Prednisolone | 21679, 2230619, 2152541, 2245532 |
|  | Methylprednisolone | 1934325, 1934333, 1934341, 30759, 30767, 36129, 30988, 2245406, 2245400, 2245408, 2245407, 2241229, 2231893, 2231894, 2231895, 2232750, 2232748, 2063727, 2063697, 2063719, 2063700, 36137, 2230210, 2230211, 30678, 30651, 30643 |
|  | Betamethasone | 2237835, 36366, 2063190, 176834, 28096, 28185 |
|  | Hydrocortisone | 888222, 888230, 888206, 888214, 30910, 30929, 872520, 872539, 878618, 878626, 30635, 30600, 30619, 30627 |
|  | Dexamathasone | 2261081, 2250055, 213624, 16462, 354309, 716715, 874582, 1977547, 664227, 2204274, 2204266, 295094, 285471, 489158, 2239534, 732893, 732885, 2260301, 2237044, 2260298, 2237046, 2237045, 1946897, 1964976, 1964968, 1964070, 2279363, 783900, 751863, 2311267, 2240687, 2240685, 2240684 |
| Other xanthines | Theophylline, combination | 545090, 476374, 334510, 356123, 792942, 721301, 317225, 828718, 640093, 828726, 828742, 307548 |
| Reliever Medications | | |
| Short-acting beta-agonists (SABA) | Salbutamol | [01926934](https://pharmacareformularysearch.gov.bc.ca/faces/Search.xhtml), [02146843](https://pharmacareformularysearch.gov.bc.ca/faces/Search.xhtml), [02146851](https://pharmacareformularysearch.gov.bc.ca/faces/Search.xhtml), [02173360](https://pharmacareformularysearch.gov.bc.ca/faces/Search.xhtml), [02208229](https://pharmacareformularysearch.gov.bc.ca/faces/Search.xhtml), 02208237, [02208245](https://pharmacareformularysearch.gov.bc.ca/faces/Search.xhtml), [02213419](https://pharmacareformularysearch.gov.bc.ca/faces/Search.xhtml), [02213427](https://pharmacareformularysearch.gov.bc.ca/faces/Search.xhtml), [02213486](https://pharmacareformularysearch.gov.bc.ca/faces/Search.xhtml), [02232570](https://pharmacareformularysearch.gov.bc.ca/faces/Search.xhtml), [02239365](https://pharmacareformularysearch.gov.bc.ca/faces/Search.xhtml), 02241497, [02243115](https://pharmacareformularysearch.gov.bc.ca/faces/Search.xhtml), [02245669](https://pharmacareformularysearch.gov.bc.ca/faces/Search.xhtml), [02326450](https://pharmacareformularysearch.gov.bc.ca/faces/Search.xhtml), 02419858 |
|  | Terbutaline | [00786616](https://pharmacareformularysearch.gov.bc.ca/faces/Search.xhtml) |
|  | Fenoterol | [00541389](https://health-products.canada.ca/dpd-bdpp/info.do?lang=en&code=4606), [02006383](https://health-products.canada.ca/dpd-bdpp/info.do?lang=en&code=13409), [00371807](https://health-products.canada.ca/dpd-bdpp/info.do?lang=en&code=2700), [00454796](https://health-products.canada.ca/dpd-bdpp/info.do?lang=en&code=3545), [02056712](https://health-products.canada.ca/dpd-bdpp/info.do?lang=en&code=16197), [02056704](https://health-products.canada.ca/dpd-bdpp/info.do?lang=en&code=16196) |
